# Linker histone H1FOO is required for bovine preimplantation development by regulating lineage specification and nucleosome assembly

**DOI:** 10.1101/2021.12.07.471683

**Authors:** Shuang Li, Yan Shi, Yanna Dang, Bingjie Hu, Lieying Xiao, Panpan Zhao, Shaohua Wang, Kun Zhang

## Abstract

Linker histone H1 binds to the nucleosome and is implicated in the regulation of the chromatin structure and function. The H1 variant H1FOO is heavily expressed in oocytes and early embryos. However, given the poor homology of H1FOO among mammals, the functional role of H1FOO during early embryonic development remains largely unknown, especially in domestic animals. Here, we find that H1FOO is not only expressed in oocytes and early embryos but granulosa cells and spermatids in cattle. We then demonstrate that the interference of H1FOO results in early embryonic developmental arrest in cattle using either RNA editing or Trim-Away approach. H1FOO depletion leads to compromised expression of critical lineage-specific genes at the morula stage and affects the establishment of cell polarity. Interestingly, H1FOO depletion causes a significant increase in expression genes encoding other linker H1 and core histones. Concurrently, there is an increase of H3K9me3 and H3K27me3, two markers of repressive chromatin and a decrease of H4K16ac, a marker of open chromatin. Importantly, overexpression of bovine H1FOO results in severe embryonic developmental defects. In sum, we propose that H1FOO controls the proper chromatin structure that is crucial for the fidelity of cell polarization and lineage specification during bovine early development.

## INTRODUCTION

As the basic chromatin unit, the nucleosome consists of an octamer of core histones (H2A, H2B, H3, and H4) that surrounded by 147 bp DNA and linker DNA with a diverse length bound to the linker histone H1 (Simpson, 1978; Syed et al., 2010). Unlike core histones, linker histones exhibit high diversity in amino acid sequence (Izzo et al., 2008; Martire and Banaszynski, 2020; Prendergast and Reinberg, 2021). There are 11 H1 variants in mammals, including seven somatic variants (H1.1 to H1.5, H1.0 and H1x), and four germline-specific variants (H1foo, H1t, H1fnt, and Hils1)(Happel and Doenecke, 2009; Martianov et al., 2005; Tanaka et al., 2001; Yan et al., 2003). Since the discovery of nucleosome, a great progress has been made as for the understanding of core histones and how their variants and chemical modifications affect chromatin structure and gene expression. However, less attention has been paid on the functional role of linker H1.

It has been well established that linker histones are generally involved in the regulation of chromatin condensation, heterochromatin formation, and gene expression (Fyodorov et al., 2018; Zhou and Bai, 2019). For example, H1 interacts directly with H3K9me3 methyltransferases and promote chromatin compaction in mouse embryonic stem cells (Bulut-Karslioglu et al., 2014; Healton et al., 2020). However, these traditional views have been challenged by recent studies (Prendergast and Reinberg, 2021).

A substitution of linker H1 variants occurs during oogenesis and early embryogenesis in a number of species. In Drosophila, histone BigH1, is switched to somatic H1 in most cells by cellularization when the zygotic genome is activated (Perez-Montero et al., 2013). Functional studies reveal an essential role of BigH1 in zygotic genome activation (ZGA). In 2001, H1FOO (also known as H1.8, homolog to BigH1) is an oocyte H1 variant that was discovered in mice. H1FOO is maternally deposited into the oocyte is later replaced with somatic H1 at the time of ZGA (Funaya et al., 2018; McGraw et al., 2006).

H1FOO plays a critical role independent of other H1 variants in early embryonic development. Histone H1c/H1d/H1e triple-mutants die during embryonic development, but individual depletion of H1c, H1d or H1e does not affect normal development in mice (Fan et al., 2003; Fan et al., 2001). In contrast to the functional redundancy of somatic H1, deletion of H1foo alone cause severe developmental defects in mouse early embryos. Although BigH1 regulates ZGA in Drosophila (Perez-Montero et al., 2013), it seems that H1Foo plays no such role in mice. It is not surprising since H1FOO are highly divergent in amino sequence among species, suggesting species-specific role of H1FOO. In cattle, we previously found that RNAi-mediated knockdown (KD) of H1FOO results in developmental arrest at the morula stage (Li et al., 2021). However, the specific functional role of H1FOO and its mechanism have yet to be determined.

Here, we explore the functional significance of H1FOO during bovine early embryonic development. We determine that H1FOO is not only present in bovine eggs and early embryos, but also exist to varying degrees in granulosa cells and testis. RNA editing and Trim-Away experiments reveal H1FOO is functionally required for bovine early development. RNA-seq analysis show the dysregulation of multiple genes involved in lineage differentiation and overrepresentation of other linker and core histone variants. In sum, our study establishes a mechanism by which the linker histone H1FOO controls critical processes that essential for bovine preimplantation development.

## RESULTS

### H1FOO is not exclusively expressed in oocytes and early cleaved embryos in cattle and humans

To explore the function of H1FOO in bovine embryos, we first made an antibody that targets the bovine H1FOO-specific sequence. RNAi-mediated silencing of H1FOO validated the specificity of the antibody (Fig. 1 A). H1FOO exhibits a typical expression pattern of a maternal protein at both mRNA and protein level (Fig. 1B and S1A). Furthermore, the intensity of H1FOO is diminished in both the cytoplasm and the nucleus (Fig. S1B and S1C). Interestingly, results of immunohistochemistry and Western Blot reveal H1FOO was expressed, despite at low level, in granulosa cells (Fig 1C, 1D, 1E). Interestingly, H1FOO was also highly expressed in spermatids of bovine testis, but not detected in sperm cells in both epididymis head and tail (Fig. 1C, S1D and S1E), suggesting H1FOO is removed during spermiogenesis. These results indicate that H1FOO is not oocyte-specific H1. Indeed, we also found H1FOO was also detectable in human granulosa cells and the abundance of *H1FOO* mRNA is declining during oogenesis (Fig S2A, S2B and S2C).

**Figure 1.**
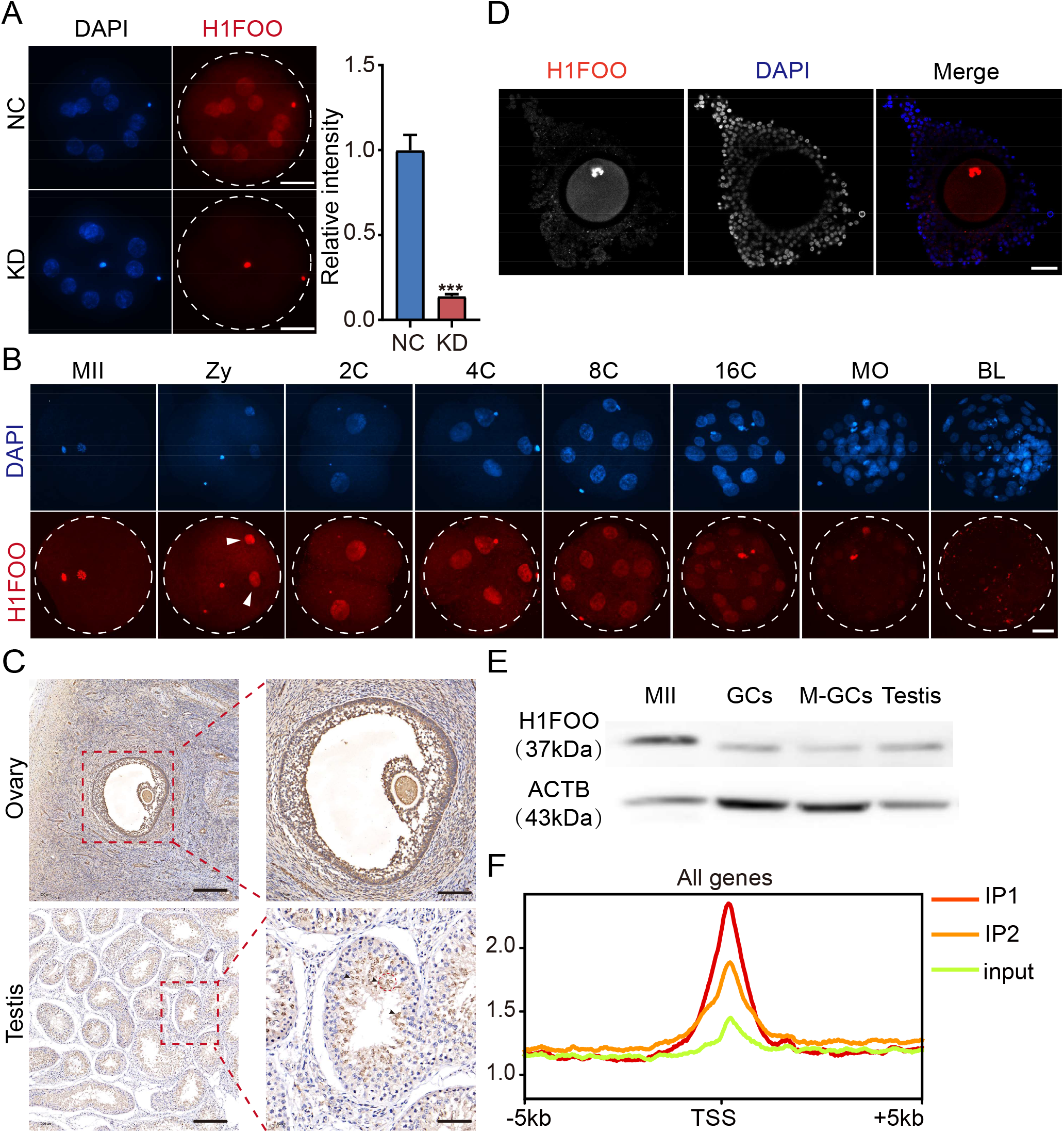
H1FOO is not exclusively expressed in oocytes and early cleaved embryos in cattle and humans. (A) Left: immunostaining validation for H1FOO knockdown (KD) efficiency at 8-cell stages (three replicates of 5-8 embryos were analyzed per group). Scale bar = 50 μM. Right: analysis of the relative intensity of H1FOO KD for experiments shown in bar charts. *\*\*\*P* < 0 .001. (B) IF detection of H1FOO protein during oocyte maturation and embryonic development in cattle. Red: H1FOO protein; Blue: DAPI (Nuclei). The experiment was independently repeated two times with 3–7 embryos were examined at each stage per time. Scale bar = 50 μM. MII: metaphase II, Zy: zygote, 2C: 2-cell, 4C: 4-cell, 8C: 8-cell, 16C: 16-cell, MO: morula, BL: blastocyst. Arrowheads indicate the pronucleus. (C) Immunohistochemistry analysis of H1FOO location in cumulus-oocyte complexes (COCs; above) and seminiferous tubule (below). Dash squares indicated the magnified regions in ovary (above) and testis (below). Arrowheads indicate the undeformed sperm cell. Scale bar = 100 μM. (D) IF detection of H1FOO protein in COCs. The single section of image shown here. Red: H1FOO protein; Blue: DAPI. Scale bar = 50 μM. (E) Western blot assay for H1FOO expression in MII, granulosa cells (GCs), mature granulosa cells (M-GCs) and testis (200 oocytes, 60-80 ug tissue samples). (F) Distribution of bovine H1FOO around the transcriptional start site (TSS) in 8-16-stage embryos (E3.0).

To further illustrate the chromatin identity of H1FOO target regions, we performed ChIP-seq to examine the distribution of H1FOO on the chromatin at 8-16-cell stage. Collectively, H1FOO was mainly enriched at the promoter regions (Fig. 1F).

### H1FOO depletion leads to developmental failure during morula to blastocyst transition in cattle

In contrast with mice (Funaya et al., 2018), we previously demonstrated that RNAi-mediated silencing of H1FOO results in a developmental arrest at morula stage in cattle (Li et al., 2021). Given the concern on the efficiency and off-target effects of RNAi approach, we further tested the functional role of H1FOO by using two other independent approaches including CRISPR-Cas13d and Trim-Away.

CRISPR-Cas13d system is an RNA editing approach that can efficiently deplete maternally-stored transcripts in early embryos (Kushawah et al., 2020). We designed three single-guide RNAs (sgRNAs) and injected them together with Cas13d mRNA into the zygote (Fig. 2A). These sgRNAs all display robust efficiency in knocking-down exogenous H1FOO in early mouse embryos (Fig. S3A to S3B). Results of injection into bovine zygotes showed that H1FOO protein abolished sharply at the 8-cell stage (Fig. 2B, Fig. S4A). Similarly, the developmental capability is greatly inhibited in Cas13d KD groups with an arrest during the morula to blastocyst transition (Fig. 2C and D).

**Figure 2.**
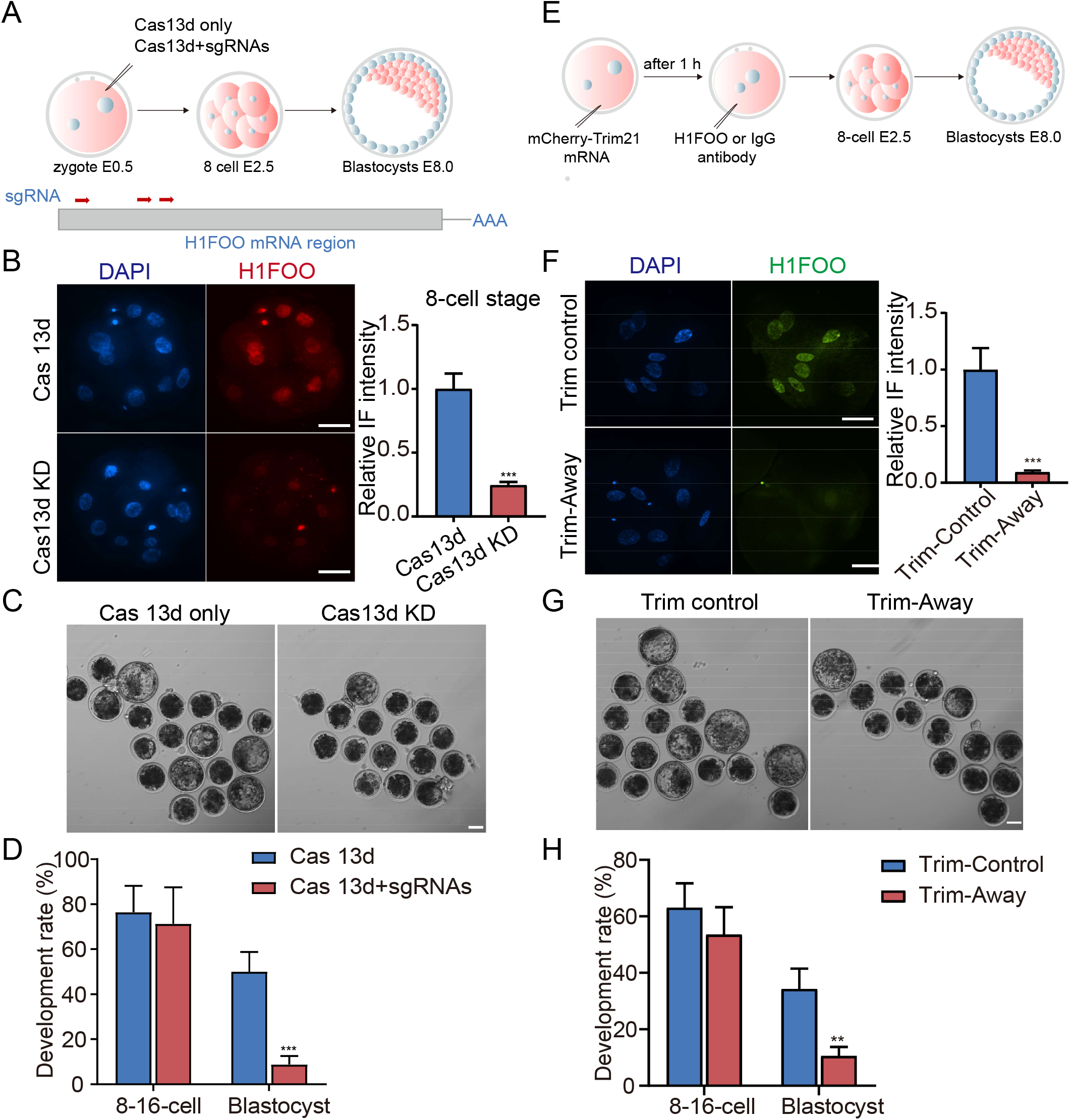
H1FOO depletion leads to developmental failure during morula to blastocyst transition in cattle. (A) Schematic of CRISPR-Cas13d strategy to deplete H1FOO mRNA. sgRNAs targeting different exon coding regions of H1FOO. (B) Immunostaining validation for H1FOO KD efficiency at 8-cell stage by CRISPR-Cas13d system. Scale bar = 50 μM. Analysis of the relative intensity of H1FOO KD for experiments shown in bar chart. n=2; 5-8 embryos were analyzed per group. *\*\*\*P* < 0 .001. (C) Representative images of bovine embryos in only Cas13d mRNA, mixture of Cas 13d KD mRNA and sgRNAs injected groups at day 8. Scale bars = 100 μm. (D) Effects of CRISPR-Cas13d induced H1FOO KD on 8-16-cell (E3.0) and blastocyst (E8.0) rates of bovine embryos. n=3 experiments; *\*\*\*P* < 0.001. (E) Schematic of Trim-Away strategy to deplete H1FOO protein. (F) IF validation for H1FOO KD efficiency at 8-cell stage by Trim-Away. Scale bar = 50 μM. Analysis of the relative intensity of H1FOO KD for experiments shown in bar chart. n=2 expreriments, 8 embryos were analyzed per group. *\*\*\*P* < 0 .001. (G) Representative images of bovine embryos in Trim control and Trim-Away groups at day 8. Scale bars = 100 μm. (H) Effects of Trim-Away induced H1FOO depletion on 8-16-cell (E3.0) and blastocyst (E8.0) rates of bovine embryos. n=3 experiments. *\*\*\*P* < 0.001.

Trim-Away approach has been used to rapidly eliminate endogenous proteins without prior modification (Clift et al., 2018). We first microinjected mCherry-TRIM21 mRNA into the fertilized eggs, and then injected H1FOO or IgG antibodies into the embryo (Fig. 2E and S4B). Remarkably, H1FOO was greatly degraded following microinjection (Fig. 2F, S4C). The blastocyst rate was also significantly decreased in the Trim-Away groups (Fig. 2G and 2H). In summary, these data indicate H1FOO is required for bovine early embryonic development, especially for the morula to blastocyst transition, and the role of H1FOO is species-specific.

### H1FOO depletion results in significant disruption of the transcriptomic profiles at morula stage

Considering the significant decline of H1FOO during maternal-to-zygotic transition, we sought to explore whether H1FOO plays a role in ZGA. To test this proposal, we firstly assessed the effect of H1FOO KD on the transcriptional activity during ZGA initiation. IF results showed that H1FOO KD caused phosphorylated Ser2 (Ser2P) of RNA polymerase II (RNAP2), active marker of transcriptional activity (Zaborowska et al., 2016), distinctly decreased at the 8-16 cell stage (Fig. 3A).

**Figure 3.**
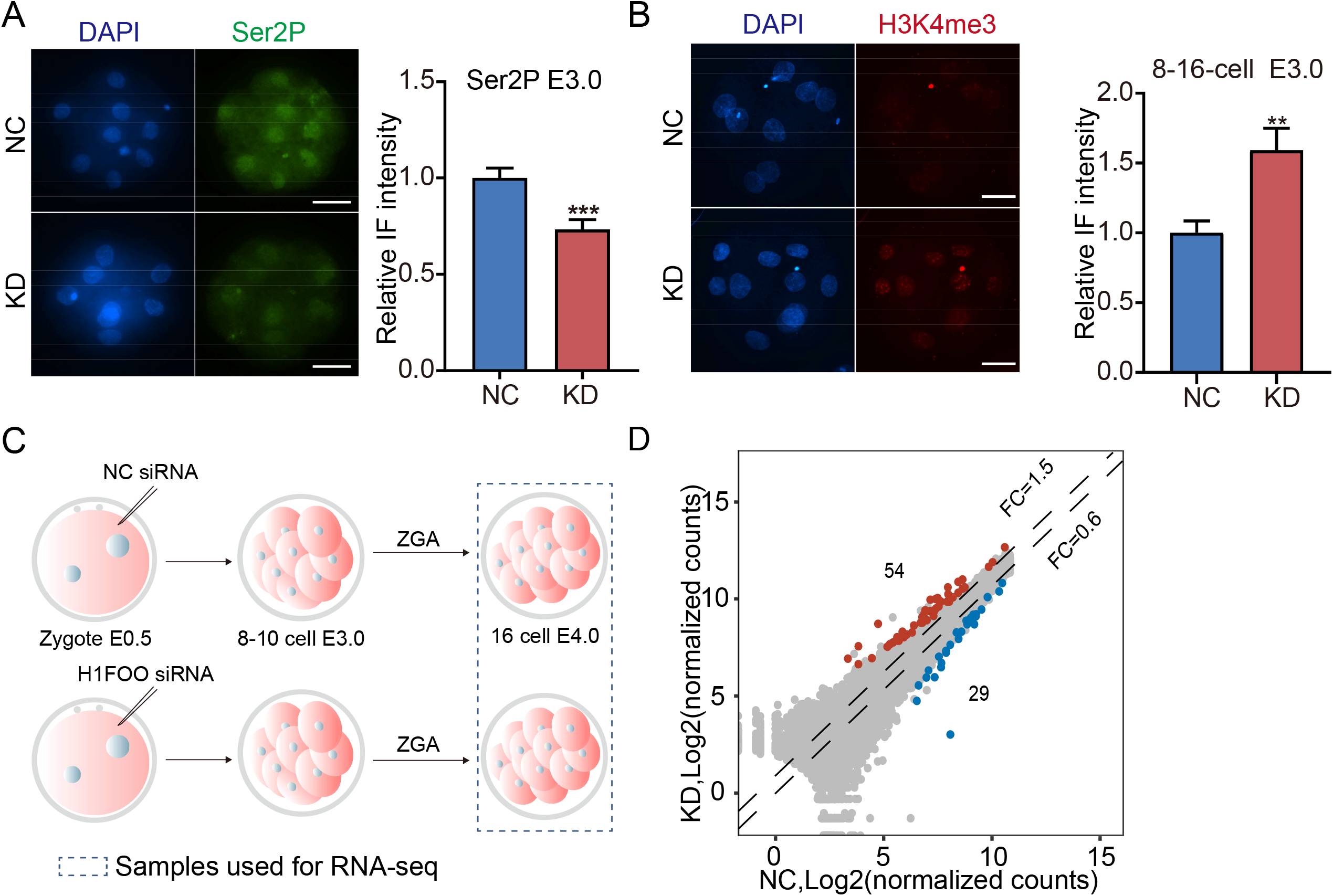
H1FOO reduction influences H3K4me3 reprogramming at 8-16-cell-stage.(A) IF analysis of phosphorylated RNA polymerase II (Ser2P) in NC and H1FOO KD embryos (E3.0). Scale bar = 50 μM. 23 embryos from three replicates were analyzed per group. Analysis of the relative intensity of H1FOO KD for experiments is shown in bar charts. *\*\*\*P* < 0 .001. (B) IF detection of H3K4me3 signal at 8-16-cell stage (E). Scale bar = 50 μM. Three independent experiments were performed and 18-22 embryos were analyzed in total. *\*\*P* < 0 .01. (C) Schematic overview of the samples collected for RNA-seq analysis (E4.0). (D) Volcano plots of all genes detected at 16-cell stage in NC and KD groups. Red dots represent upregulated genes (FC > 1.5, *P* adjusted < 0.05) and blue dots downregulated genes (FC < 0.6, P adjusted < 0.05).

Histone H3 lysine 4 trimethylation (H3K4me3) is a hallmark of active genes and decreased during ZGA (Dahl et al., 2016; Zhou et al., 2020). We observed the significant disappearance of H3K4me3 at the 8-16 cell stage in NC not in KD embryos (Fig. 3B). These data suggest H1FOO loss already leads to abnormal epigenetic reprograming at 8-16 cell stage.

However, RNA-seq result shows that only 83 differentially expressed genes (DEGs) were found in KD compared with NC group (FC >= 1.5 or <=0.6; Padj <=0.05), including 54 up-regulated genes and 29 down-regulated genes (Fig. 3C, 3D and S5).

Since most embryos were arrested at the morula stage in KD group, we next examined the molecular outcome of H1FOO loss at the morula stage (Fig. 4A). Samples were collected when there is no significant difference in total cell number per embryo (Fig. 4B). Results showed 837 transcripts were altered, of which 338 were up-regulated and 449 were down-regulated (Fig. 4C). Association analysis of DEGs showed that only 14 transcripts commonly differed both in the 16-cell and morula stages (Fig. 4D).

**Figure 4.**
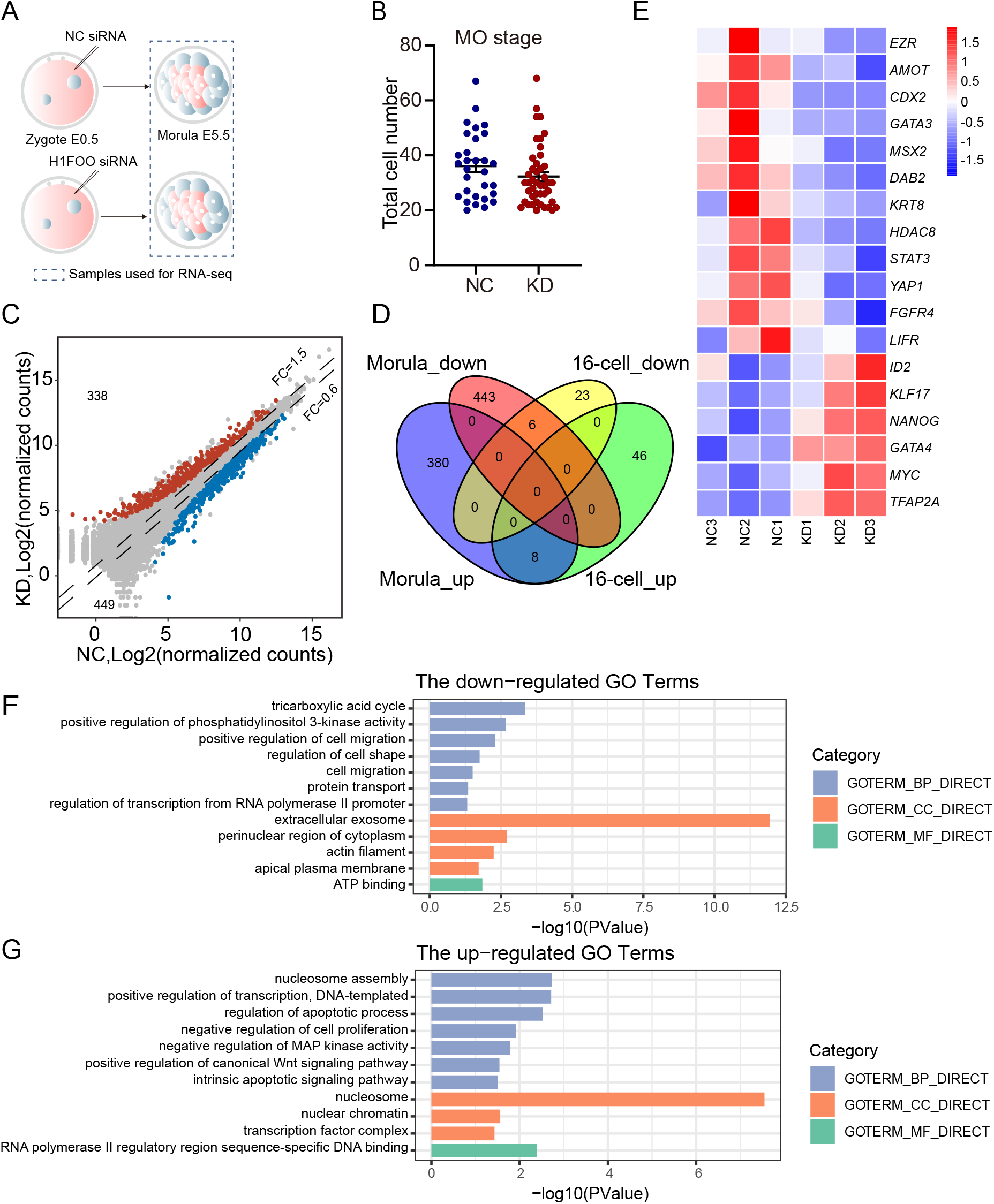
H1FOO deficiency causes a large-scale disruption of the transcriptome profile in bovine morula. (A) Schematic overview of the samples at morula stage collected for RNA-seq analysis. (B) Validation of total cell number per day 5.5 morula in NC and KD groups. Three independent experiments were performed and each embryo was stained with DAPI to confirm cell number. Each dot represents an embryo. (C) Volcano plots of all genes detected at MO stage in NC and KD groups. Red dots represent upregulated genes (FC > 1.5, *P* adjusted < 0.05) and blue dots downregulated genes (FC < 0.6, P adjusted < 0.05). (D) Venn diagram illustrating the common differentially expressed genes at 16-cell and MO stage identified in KD embryos relative to NC groups. (E) Overrepresentation of genes related to lineage specification among DEGs. (F) Gene ontology (GO) analysis of downregulated genes identified in KD morulaes relative to NC groups. (G) GO terms of upregulated genes identified in KD morulaes relative to NC groups.

GO analysis of down-regulated genes revealed overrepresentation of genes involved in positive regulation of cell migration and apical plasma membrane (Fig. 4E and 4F). In particular, we found genes related to impaired lineage differentiation, including *CDX2, GATA3* and *KRT8*, which are trophectoderm (TE)-specific markers (Gerri et al., 2020; Negron-Perez et al., 2017; Wei et al., 2017) and genes associated with cell polarity, including *AMOT* (Hirate et al., 2013) and *EZR* (Louvet et al., 1996), were also significantly reduced (Fig. 4E). Moreover, the most strikingly enriched GO terms of up-regulated genes are involved in nucleosome assembly, in addition of apoptotic process and negative regulation of cell proliferation (Fig. 4G). These results indicated that H1FOO-deficient morulae had already deviated from NC morulae at the molecular level even they were morphologically indistinguishable. It also suggests that H1FOO acts not only as a linker histone but regulates early lineage specification during preimplantation development.

### H1FOO KD results in compromised linage specification in day 5.5 morulae

The first lineage specification takes place during the morula-to-blastocyst transition and gives rise to ICM and TE (Rossant, 2018). Because we noted down-regulated expression of key lineage-specific genes among DEGs from RNA-seq analyses (Fig. 3F), we next tested if the early differentiation program was abnormal in KD morulae. IF assays revealed that the amounts of CDX2 and GATA3 were all reduced in KD morulae (Fig. 5A). HDAC8 is abundantly expressed in the ICM (Negron-Perez et al., 2017). We detected a significant decrease in HDAC8 protein of KD embryos (Fig. 5B), which is consistent with RNA-seq results. These data suggest a failure of the first lineage specification in H1FOO-deficient embryos.

**Figure 5.**
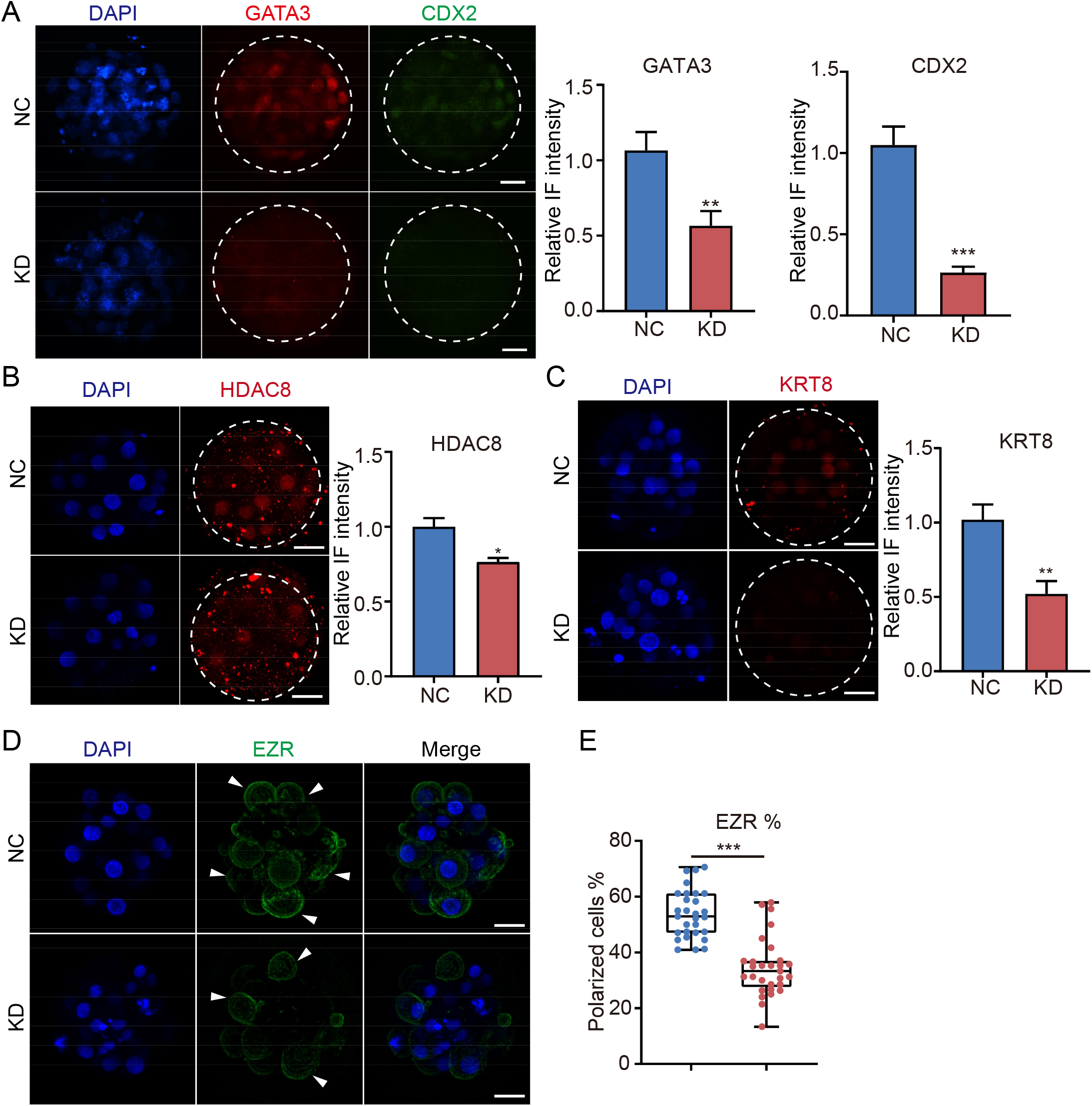
H1FOO KD results in a compromised linage specification at morula stage. (A) IF analysis of GATA3 (red) and CDX2 (green) in NC and H1FOO KD morulae. Scale bar = 50 μM. Quantification of GATA3 and CDX2 staining in the IF images by bar graphs. Three biological replicates with 6-8 embryos were analyzed per group each time. *\*\*P* < 0 .01. *\*\*\*P* < 0 .001. (B) IF detection of HDAC8 in NC and H1FOO KD morulae. Scale bar = 50 μM. Three biological replicates with total 20-23 embryos analyzed per group. *\*P* < 0 .05. (C) IF detection of KRT8 in NC and H1FOO KD morulae. Scale bar = 50 μM. Analysis of the relative intensity of KRT8 for experiments shown in bar chart. n=3. 6-8 embryos were analyzed per group each time. *\*\*P* < 0 .01. (D) IF detection of EZR in NC and H1FOO KD morulae. Scale bar = 50 μM. (E) Proportions of polarized cell (EZR^+^) in morulae. Each dot represents an embryo. *\*\*\*P* < 0.001. n=4. 7-10 embryos were analyzed per group each time.

Apical domain formation is an essential process for embryo polarization to induce the segregation of the ICM and TE lineages (Korotkevich et al., 2017). In this process, the ERM proteins (Ezr, Radixin, and Moesin) and Par protein complex surrounded by an actomyosin ring migrate to the apical region of polar cells in mice (Plusa et al., 2005; Zhu et al., 2020). Hippo pathway is involved in the regulation of the expression of important polar proteins (Rossant, 2018). We found that Keratin8 (KRT8) became enriched at the apical regions of the outer cells along with development (Fig. S6A), and it decreased substantially in H1FOO KD nucleus (Fig. 5C). Meanwhile, EZR was specifically expressed at the apical membrane of outer cells in the bovine (Fig. 5D). However, the number of polarized cells (EZR^+^) is greatly reduced in KD embryos (Fig. 5D and 5E). However, we found YAP1 signal is normal in KD morula (Fig. S6B). Altogether, these data suggest that H1FOO regulates the first lineage specification event likely through regulation of apical domain establishment.

### Chromatin composition and structure are disturbed in H1FOO-deficient embryos

H1 plays an important role in maintaining chromatin configuration and stability (Climent-Canto et al., 2020; Funaya et al., 2018; Henn et al., 2020). Surprisingly, H1FOO depletion results in elevated expression of multiple genes encoding linker histone and core histone variants (Fig. 6A and S7A). IF results confirmed H2A protein was increased in KD morulae (Fig. 6B).

**Figure 6.**
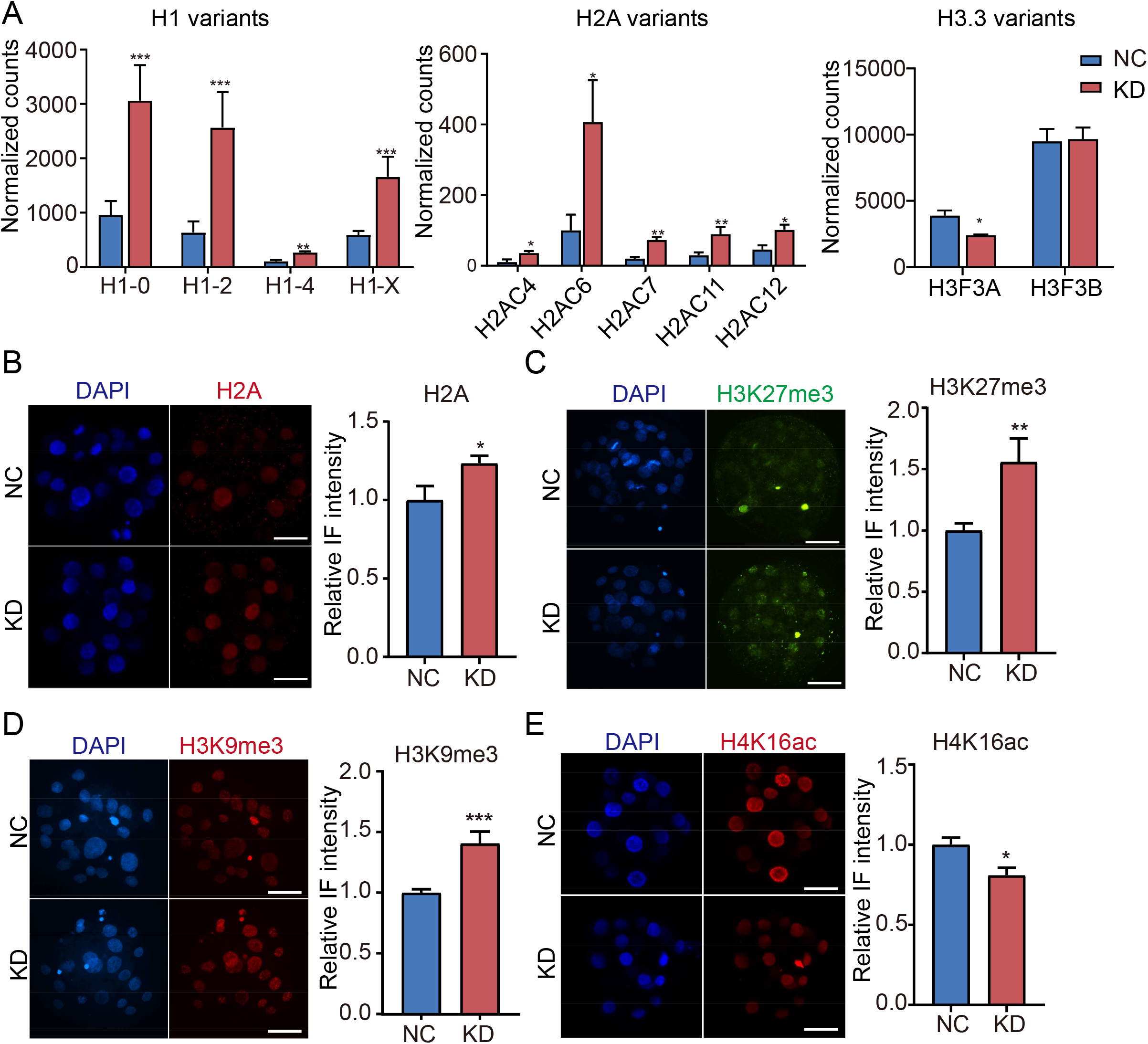
Histone methylation and acetylation were perturbed in H1FOO deficient embryos with increased histone variants. (A) RNA-seq results related to the expression levels of histone H1, H2A and H3.3 variants in day 5.5 morulaes. *\*P* adjusted < 0.05, \**\*P* adjusted < 0.01, *\*\*\*P* adjusted < 0.001. (B) Immunostaining validation of core histone H2A in morulae. Scale bar = 50 μM. Quantification of H2A staining in the IF images by bar graph. n=3. 5-8 embryos were analyzed per group each time. *\*P* < 0.05. (C and D) IF staining of repressive histone markers H3K27me3 (C) and H3K9me3(D) in NC and KD morulae (E5.5). Scale bar = 50 μM. Analysis of the relative intensity of H3K27me3 and H3K9me3 for experiments shown in bar charts. \**\*P* < 0.01, *\*\*\*P* < 0 .001. n=3. 6-8 embryos were analyzed per group each time. (E) IF validation of H4K16ac in morulae. Scale bar = 50 μM. Quantification of H4K16ac staining in the IF images by bar graph. n=2 experiment. 8-10 embryos were analyzed per group each time. *\*P* < 0.05.

H3K9me3 and H3K27me3 signals markedly increased in the KD nucleus compared with NC (Fig. 6C and 6D)(Fyodorov et al., 2018). Moreover, we found that the intensity of H4K16ac decreased notably while H3K36me2 showed no significant difference (Fig. 6E and S7B). Collectively, these results indicate that disordered histone modifications and increased nucleosome assembly may be responsible for the dysregulation of the transcriptome in H1FOO KD embryos.

### Overexpression of bovine H1FOO impairs the developmental potential of early embryonic embryos in cattle and mice

Next, we wondered if overexpression of H1FOO affects bovine early embryonic development. Thus, we microinjected H1FOO mRNA tagged with FLAG into bovine zygotes and found that the level of H1FOO was significantly increased (Fig. 7A and 7B). Surprisingly, the proportion of embryos to become 8-16-cell embryos is decreased in OE groups (P<0.05), while the blastocyst rate is slightly decreased (p=0.22; Fig. 7C). Meanwhile, both total cell number (DAPI^+^) and TE cell number (CDX2^+^) per blastocyst were decreased significantly (P<0.05) whereas the number of ICM (SOX2^+^) did not change (Fig. 7D and 7E). To understand whether H1FOO OE promotes heterochromatin formation as somatic H1 (Fyodorov et al., 2018), we detected the level of H3K9me3 and found no difference between NC and OE group (Fig. 7F). This result demonstrated the H1FOO behave in a different manner with somatic H1.

**Figure 7.**
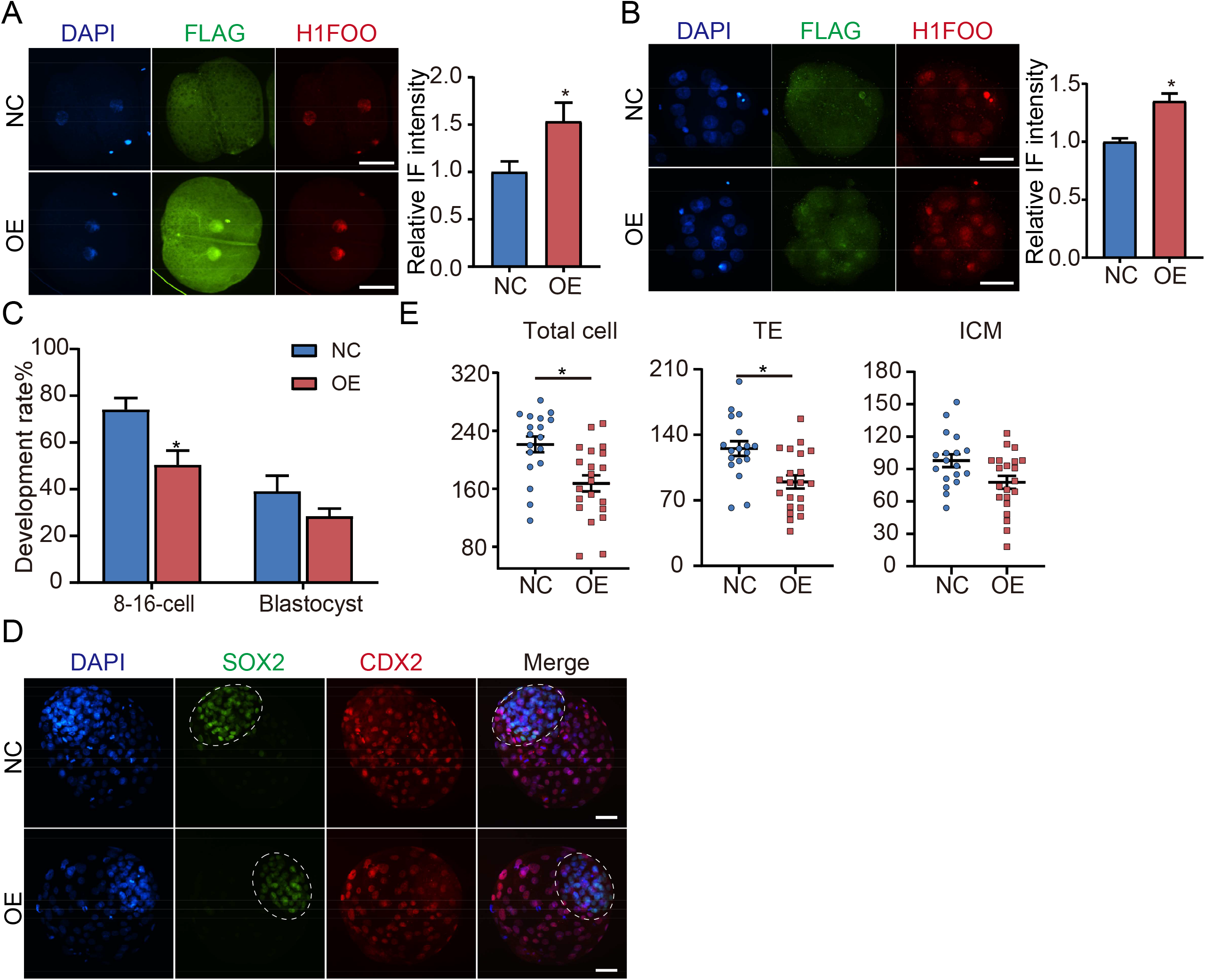
Overexpression of H1FOO impairs the developmental potential of bovine early embryos. (A and B) IF staining was performed to analyze the level of overexpressed H1FOO mRNA at 2-cell (A) and 16-cell (B) stages. Green: FLAG protein represents exogenous H1FOO. Red: endogenous and exogenous H1FOO protein. Scale bar = 50 μM. Analysis of the relative intensity of H1FOO for experiments shown in bar charts. n=2 experiments, 12-15 embryos were analyzed per group. *\*P* < 0 .05. (C) Effects of bovine H1FOO OE on 8-16-cell (E3.0) and blastocyst (E8.0) rates of bovine embryos. n=3 experiments. *\*P* < 0.05. (D) Representative IF pictures of SOX2 (red) and CDX2 (green) in NC and H1FOO OE groups at blastocyst stage (E8.0). The nuclei were labeled by DAPI. The circle indicates the ICM. Scale bar = 50 μM. (E) The number of total cells (DAPI^+^), TE cells (CDX2^+^), and ICM cells (SOX2^+^) per blastocyst in NC and H1FOO OE groups(E8.0). n=3 replicates, 6-8 blastocysts per group each time. *\*P* < 0 .05.

To compare the functional role of H1FOO between species, we overexpressed mouse and bovine H1FOO in mouse preimplantation embryos. Results display a significant OE efficiency (Fig. 8B and 8C). In vitro culture of embryos shows that OE of bovine H1FOO (bH1FOO) in early mouse embryos resulted in severe developmental disorders, but OE of mouse H1foo mRNA had no effect (Fig. 8D and 8E), suggesting H1FOO’s role is species-dependent. Thus, these results suggest that H1FOO OE is not conducive to blastocyst development in both cattle and mice.

**Figure 8.**
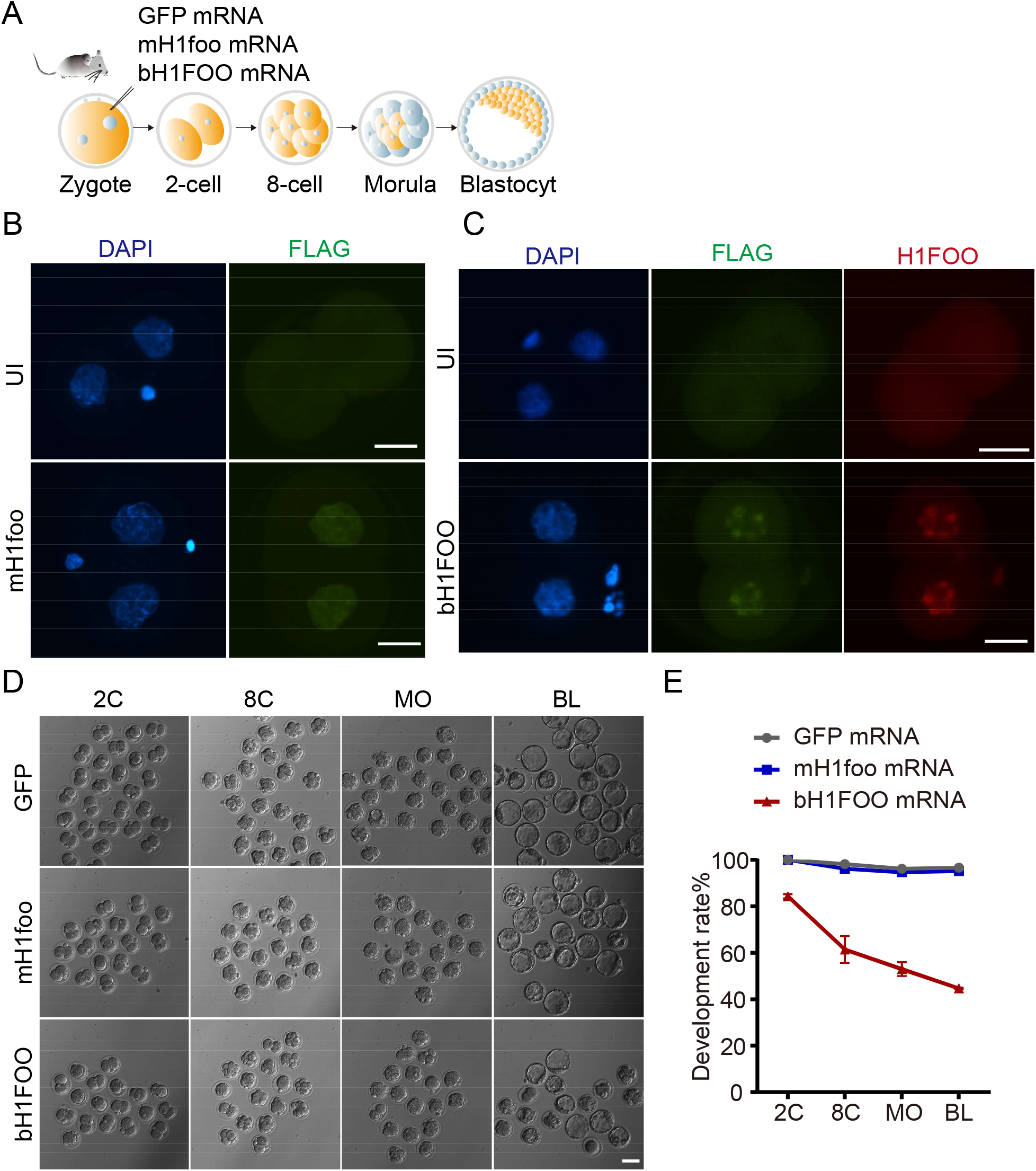
Overexpression of bovine H1FOO impairs the developmental potential of mouse embryos. (A) Schematic of H1FOO mRNA overexpression (OE) strategy to detect the development rates in mouse embryos. The fertilized eggs of the mice were divided into three groups and injected with bovine H1FOO mRNA (bH1FOO), mouse H1foo (mH1foo) mRNA and GFP mRNA respectively. (B) IF detection for OE level of mH1foo tagged with FLAG (green) at mouse 2-cell stage (about 20 hours post-injection). Scale bar = 50 μM. (C) IF detection for OE level of bH1FOO (red) tagged with FLAG (green) at mouse 2-cell stage (about 20 hours post-injection). Scale bar = 50 μM. (D) Representative images of mouse embryos injected with different Mrna Nduring preimplantation development. Scale bars = 100 μm. (E) Effects of bH1FOO and mH1foo OE on developmental rates of mouse embryos. n=2.

## DISCUSSION

We report here that the linker histone variant H1FOO is necessary for bovine early embryonic development using three independent approaches. H1FOO deficiency leads to a significant disruption to the transcriptome. Importantly, H1FOO deficiency is detrimental to the first lineage specification event. Moreover, H1FOO is required to maintain the expression profiles of core and linker histones, and essential for fidelity of histone modifications in bovine early embryos (Fig. 9).

**Figure 9.**
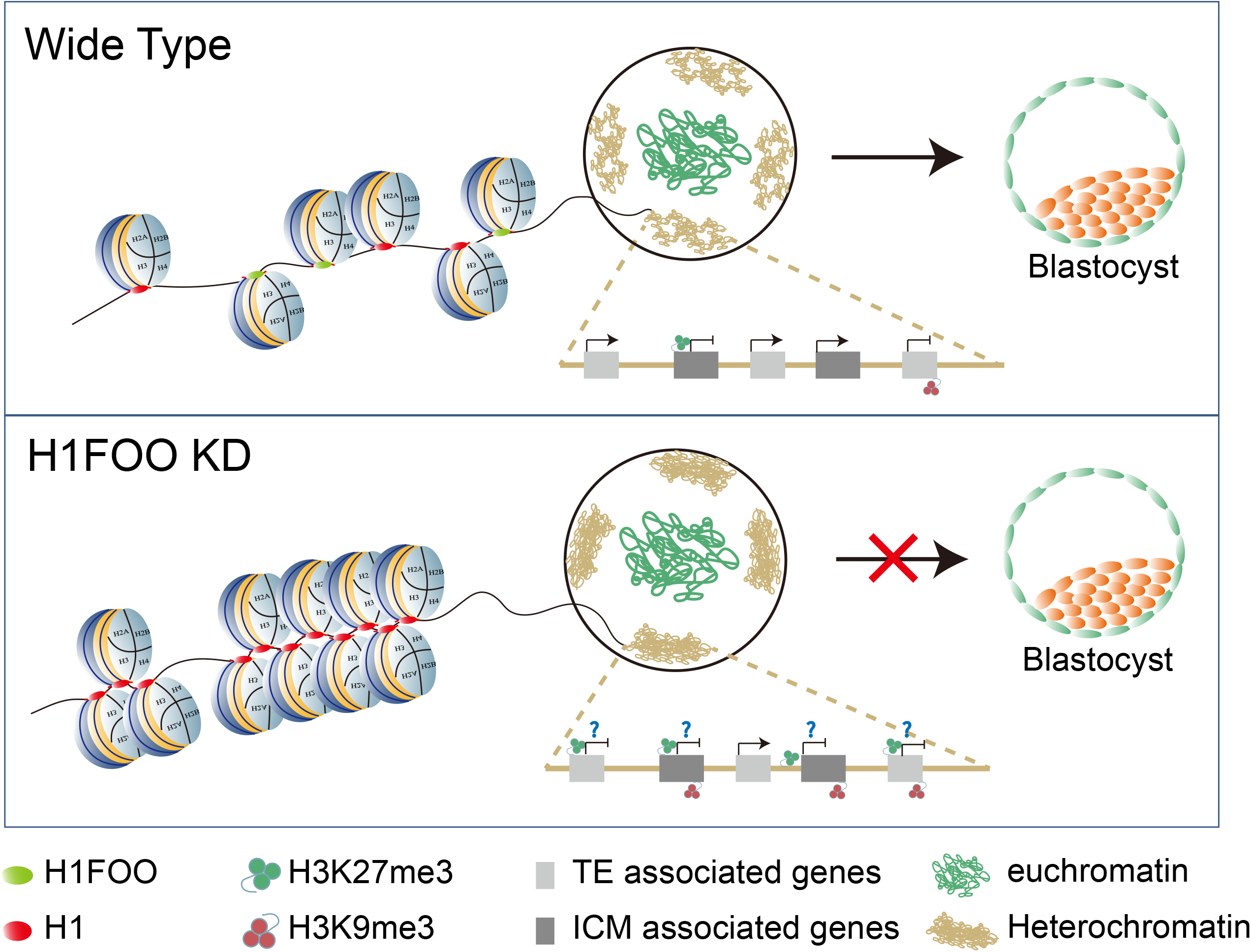
Working model of how H1FOO is proposed to facilitate lineage specification and chromatin structure in bovine early embryos. In the wild-type embryos (top), H1FOO acts to maintain normal nucleosome density, especially in the heterochromatin region, to ensure correct TE and ICM associated gene expression. In the absence of H1FOO (bottom), the nucleosomes are positioned more densely with an overall increase in some heterochromatic markers, such as H3K9me3 and H3K27me3. The expression of many differentiation-related marker genes (CDX2, GATA3, and KRT8 etc.) are impaired due to changes in chromatin status. Each large oval represents a nucleus at morula stage.

Somatic linker histones mainly promote transcription silencing by regulating histone acetylation or methylation (Herrera et al., 2000; Sun et al., 2015; Willcockson et al., 2021; Yusufova et al., 2021). However, it appears that H1FOO plays an opposite function to somatic H1 in chromatin remodeling, histone epigenetic modifications and transcriptional regulation (Hayakawa and Tanaka, 2021). Our results indicated that H1FOO prefers to bind the open chromatin region at 8-16-cell stage, and the deletion of H1FOO increased histone modifications related to heterochromatin formation (H3K9me3 and H3K27me3). Interestingly, H3K27me3 is preferentially occupying the promoters of developmental genes, and early lineage specification is accompanied by asymmetric H3K27me3 enrichment in TE and ICM cell (Schwartz and Pirrotta, 2007; Xia et al., 2019). H3K9me3 also regulates cell fate transition between totipotency and pluripotent state in mouse 2C-like cell (Wu et al., 2020). Abnormal elevation of these two repressive histone marks may contribute to the impairment of the first lineage differentiation in KD embryos (Fig. 9).

H1FOO (H1s) is found in oocytes and early embryos of various mammalian and non-mammalian species (such as Xenopus, Drosophila, and sea urchin), and contributes to the stability of chromatin structure among them. However, the functional role of H1Foo varies among species. In Drosophila, BigH1 ensures transcriptional activation by controlling the rapid nucleosome reassembly at the initial stage of embryogenesis (Climent-Canto et al., 2020; Henn et al., 2020). H1foo in mice is required to form a loose chromatin structure, and its depletion delays the timing of cleavage into the two-cell stage and increases deposition of the histone H3 variant (H3.1/3.2) in one-cell stage embryos (Funaya et al., 2018). However, we found that H1FOO depletion results in embryonic arrest at the morula stage (Li et al., 2021). These results indicate that H1FOO’s role is species-dependent.

Linker histones possess the typical tripartite structure of variant proteins (Allan et al., 1980), including a globular structure, N- and C-terminal regions. These three domains have different affinities in nucleosome assembly, chromatin folding, and interaction with histone modifying enzymes (Caterino and Hayes, 2011; Vyas and Brown, 2012; Zhou et al., 2013). For example, C-terminal domain (CTD) of histone H1d is required for its physical and functional interactions with DNA and histone methyltransferases to DNA methylation and histone H3 methylation (Healton et al., 2020; Yang et al., 2013). However, the sequence similarity of proteins is low between H1FOO and somatic H1 variants, especially CTD (Fyodorov et al., 2018), thus H1FOO’s functional specificity may be attributed to the C-terminal domain.

In summary, our study firstly established CRISP-Cas13d and Trim-Away systems to explore the role of maternal H1FOO in early bovine embryo development. Our data show that the developmental arrest upon H1FOO deletion could be mainly accounted for by the impaired cell polarity and lineage differentiation. Moreover, H1FOO participates in regulating the stability of nucleosome assembly and chromatin modifications in early embryos.

## MATERIALS AND METHODS

Unless otherwise stated, reagents and chemicals were commercially obtained from Sigma-Aldrich (St. Louis, MO).

### Bovine oocyte and embryo production in vitro

Bovine embryo production in vitro were performed according to procedures as published previously (Li et al., 2021). Briefly, bovine ovaries were obtained at a local slaughterhouse. Cumulus oocyte complexes (COCs) containing more than three layers of cumulus cells were retrieved from 3–8-mm follicles at the surface of bovine ovaries. COCs were matured in Medium-199 (M4530) supplemented with 10% FBS (Gibco-BRL), 1 IU/ml FSH (Sansheng Biological Technology), 0.1 IU/ml LH (Solabio), 1 mM Sodium Pyruvate (Thermo Fisher Scientific), 2.5 mM GlutaMAX™ (Thermo Fisher Scientific), and 10 μg/ml Gentamicin at 38.5°C under 5% CO2 in humidified air for 22-24 h. Upon maturation, COCs (60∼100 COCs per well in 4-well plates) were co-incubated with spermatozoa (1∼5×106) that purified from frozen-thawed semen by a Percoll gradient. IVF was performed at 38.5°C under 5% CO_2_ for 9-12 h. Granulosa cells (GCs) were removed from the oocytes by pipetting up and down with 1 mg/ml hyaluronidase. Embryos were cultured in BO-IVC medium (IVF bioscience) for 8 days. The embryos that cleaved to 8-16 cell and became blastocysts were assessed at embryonic day 3 (E3.0) and day 8 (E8.0) after fertilization, respectively.

### Mouse H1foo mRNA synthesis in vitro

The wild-type mouse *H1foo* mRNA tagged with FLAG was constructed as before (Li et al., 2021). The coding sequence of wild-type mouse *H1foo* was amplified from cDNA libraries constructed from mouse germinal vesicle oocytes with one copy FLAG added to the 3’ end. The primer sequences are shown in Supplemental Table 1. The amplicon was subsequently cloned into a T3-driven vector. To produce complementary RNA (cRNA) in vitro, the plasmid constructed above was linearized and transcribed in vitro, capped, DNAse-treated and poly(A)-tailed using mMessage mMachine T3 Ultra Kit (Thermo Fisher Scientific). cRNAs were extracted and purified by using Mega-Clear Kit (Thermo Fisher Scientific).

### *Cas13d* mRNA and single guide RNA synthesis

pET-28b-RfxCas13d-His (Addgene #141322) plasmid containing the T7 promoter was linearized by using NotI restriction site. The linearized plasmid was transcribed in vitro based on the procedures of mMessage mMachine T7 Ultra Kit (Thermo Fisher Scientific). DNAse-treated and poly(A)-tailed *Cas13d* mRNA was purified by using Mega-Clear Kit (Thermo Fisher Scientific).

Single guide RNA (sgRNA) was designed as published previously (Kushawah et al., 2020). DNA template to generate sgRNA was generated by fill-in PCR according to procedures as published previously with slight modifications. A sgRNA universal primer containing the T7 promoter and the Cas13d component was used in combination with a sequence-specific oligo. All primer sequences are shown in Table S2. T7 in vitro transcription reaction was performed using MEGAshortscript™ T7 kit (Thermo Fisher Scientific). sgRNAs were precipitated with Sodium Acetate/Ethanol.

### Trim-Away experiment

pSMPP-mCherry-hTRIM21 vector was purchased from Addgene (#104972). mCherry-hTRIM21 was amplified using primers T7-mCherry-F and TRIM21-R (Table S2), and then product was cloned into pMD18T vector to produce in vitro-transcribed mCherry-TRIM21 mRNA. mRNA was aliquoted at a concentration of 800 ng/ul and stored at - 80°C. Rabbit anti-H1FOO primary antibody (Homemade, HuaBio, Hangzhou, China) was dissolved in 1×Phosphate Buffer Saline (PBS). Rabbit IgG primary antibody (HA1002, HuaBio) was used as Trim-control. mCherry-hTRIM21 mRNA and antibodies were microinjected separately into the embryos. mCherry fluorescence was measured 48h and 5 day after injection to validate the efficacy.

### Microinjection

*H1FOO* siRNA (25 μM) or mRNA (800 ng/μl) was microinjected in a volume of 20 pl into putative zygotes collected at 12-16h post insemination (hpi) with an inverted microscope (Nikon) equipped with a micromanipulator (Narishige). For Cas13d and sgRNAs injections, reagents were prepared and delivered on ice at the desired concentrations. Final concentration of injection mRNA mixture was 50ng/ul Cas13d and 100 ng/ul per sgRNA. For Trim-Away experiment, the fertilized eggs were first injected with 400 ng/ul *mCherry-hTRIM21* mRNA, and about 1h later, the embryos were further microinjected with 1012 ng/μl rabbit anti-H1FOO or IgG primary antibody again. To specifically detect the endogenous depletion of H1FOO, we detected H1FOO expression in the 8-cell (E2.5) and morula (E5.5) stages by immunofluorescence.

### Immunofluorescence (IF)

Samples were briefly washed with 0.1% polyvinylpyrrolidone/PBS three times, fixed with 4% paraformaldehyde/PBS for 30 minutes, and permeabilized with 0.5% Triton X-100/PBS for 30 minutes at room temperature (RT). Blocking was performed for 1h in the blocking buffer (PBS containing 10% FBS and 0.1% Triton X-100). Then, samples were incubated with primary antibodies in blocking buffer under 4°C overnight and secondary antibodies for 2h. Finally, samples were treated with 4,6-diamidino-2-phenylindoley (DAPI; Life Technologies) for 30 minutes. Images were captured with a 40× objective using an inverted epi-fluorescent microscope (Nikon, Chiyoda, Japan) or a Zeiss LSM880 confocal microscope system (Zeiss, Oberkochen, Germany). For confocal microscopy, Z-stacks were imaged with 5 μm intervals between optical sections. Stacks were projected by maximum intensity to show signals of all blastomeres in one image. Information about all antibodies is presented in Supplemental Table1

ImageJ was used to visualize images, count cell numbers, and measure signal intensity. Every primary antibody was validated to be react with bovine cells and tested in three replicates using the same microscope. Samples stained without primary antibodies was used as negative controls to verify the specificity of the antibodies used here. Depending on the experiment, the signal intensity was determined and the background was subtracted to analyze the absolute intensity.

### RNA-seq library construction and bioinformatics

Bovine embryos were harvested at 16-cell stage (E4.0; n=2; 30 embryos/group/replicate) and morula stage (E5.5; n=3; 15 embryos/group/replicate). Total RNA was extracted with a PicoPure RNA Isolation Kit. mRNA separation was achieved using oligo (dT)25 beads. Sequencing libraries were constructed with NEBNext Ultra RNA Library Prep Kit for Illumina (New England Biolabs). The libraries were sequenced with Illumina Novaseq by Novogene Co. Ltd. The raw data were trimmed with Trimmomatic (v0.36) to remove adapter sequences and low quality (q<20) bases. 15-20 million clean reads were obtained to map to UCD1.2. Then the raw counts of genes were generated by FeatureCounts, and normalized to FPKM by cufflinks. Differential expressed genes were identified by using DESeq2 package (padj ≤ 0.05 and fold change (FC) ≥ 1.5 or ≤ 0.6). Heatmaps were generated by using the pheatmap package in R. Gene ontology (GO) and Kyoto encyclopedia of genes and genomes (KEGG) analyses of differentially expressed genes were carried out by the Database for Annotation, Visualization and Integrated Discovery (DAVID).

### ULI-NChIP-seq library preparation and data processing

Embryos at 8/16-cell (E3.0) were collected (N=2 replicates, 50 embryos per group) and incubated in 0.5% pronase E for several minutes to remove the zona pellucida, and washed 3 times in 0.5% bovine serum albumin (Millpore, Billerica, MA) in DPBS (Gibco-BRL, Grand Island, NY) before flash freeze in liquid nitrogen. ULI-NChIP was performed according to previous procedure (Brind’Amour et al., 2015). One microgram of H1FOO antibody was used for each immunoprecipitation reaction. ULI-NChIP libraries were generated using the NEB Ultra DNA Library Prep Kit (E7645). The paired-end 150 bp sequencing was performed on a NovaSeq platform.

The raw sequencing reads were trimmed with Trimmomatic (version 0.39) to remove residual adapter sequences and low-quality bases. Then the clean reads were aligned to Bos Taurus UMD3.1.1 using Bowtie2 (version 2.3.5) (Langmead and Salzberg, 2012) with the default parameters. Alignments with low quality were removed by SAMtools (version 1.7)(Li et al., 2009), and PCR duplicates were removed with Picard (version 2.23). The signal of H1FOO was calculated with computeMatix from DeepTools (Ramirez et al., 2014).

### Western blot

Samples (100 oocytes, mature or immature GCs, testis) were treated with RIPA lysis buffer (Beyotime) containing phenylmethylsulfonyl fluoride (1 mM; Beyotime) on ice. Total protein in loading buffer was separated by 10% SDS-PAGE and then transferred to polyvinylidene difluoride (PVDF) membranes. Membrane was blocked for 1h in nonfat milk (5%), followed by incubation with primary antibodies overnight at 4°C and secondary antibodies for 1.5 h at RT. Signals were detected and imaged with WESTAR NOVA 2.0 (Cyanagen, Bologna, Italy). Antibody information is shown in Supplemental Table 1.

### Immunohistochemistry (IHC)

Fresh tissues (ovary, testis, head of epididymis, caudal epididymis) were excised and fixed in 4% PFA and dehydrated overnight in 70% ethanol. The fixed specimens were embedded in paraffin, cut into 5μm-thick sections and stained with hematoxylin using a standard protocol. Briefly, Sections were labeled with rabbit anti-H1FOO in a humid chamber overnight at 4°C before the use of corresponding secondary antibody (HRP) at RT for 50 minutes. Then, sections were subsequently counterstained with hematoxylin stain solution for 3 minutes at RT. Samples were observed by conventional light microscopy. Negative controls were made by replacing the primary antibody with PBS.

## Statistical analysis

All experiments were replicated at least three times unless stated. Two-tailed unpaired Student t-tests were used to compare differences between two groups with SPSS statistics (IBM, USA). One-way analysis of variance (ANOVA) was employed to determine significant differences between groups for the analysis of H1FOO protein profiles and KD efficiency followed by the Tukey’s multiple comparisons test. The intensity of fluorescence was analyzed using Image J as described above, and the intensity data were normalized to the relative channels in control groups. The graphs were constructed by GraphPad Prism 7.0 (GraphPad Soft- ware, USA). P < 0.05 indicated the data are statistically significant. Numerical values are presented as means ± SEM.

## Acknowledgments

We thank all members of the K. Zhang laboratories for their helpful discussions. This work was funded by National Natural Science Foundation of China (No. 31872348, No. 31672416, and No. 32072731 to K.Z.; No.31941007 to S.W.), Zhejiang Provincial Natural Science Foundation (LZ21C170001 to K.Z.).

